# UNIQmin, an alignment-free tool to study viral sequence diversity across taxonomic lineages: a case study of monkeypox virus

**DOI:** 10.1101/2022.08.09.503271

**Authors:** Li Chuin Chong, Asif M. Khan

## Abstract

Sequence changes in viral genomes generate protein sequence diversity that enable viruses to evade the host immune system, hindering the development of effective preventive and therapeutic interventions. Massive proliferation of sequence data provides unprecedented opportunities to study viral adaptation and evolution. Alignment-free approach removes various restrictions, otherwise posed by an alignment-dependent approach for the study of sequence diversity. The publicly available tool, UNIQmin offers an alignment-free approach for the study of viral sequence diversity at any given rank of taxonomy lineage and is big data ready. The tool performs an exhaustive search to determine the minimal set of sequences required to capture the peptidome diversity within a given dataset. This compression is possible through the removal of identical sequences and unique sequences that do not contribute effectively to the peptidome diversity pool. Herein, we describe a detailed three-part protocol utilizing UNIQmin to generate the minimal set for the purpose of viral diversity analyses, alignment-free at any rank of the taxonomy lineage, using the latest global public health threat monkeypox virus (MPX) as a case study. This protocol enables systematic diversity study across the taxonomic lineage, which are much needed for our future preparedness of a viral epidemic, in particular when data is in abundance, freely available, and alignment is not an option.

## Introduction

Infectious diseases have caused irreversible losses to human lives, contributing greatly to the global disease burden. Amongst pathogenic agents, viruses are abundant and ubiquitous, responsible for the surge in global death toll from infectious diseases. The current coronavirus disease (COVID-19) pandemic is a testimony to this, affecting more than 190 countries/geographical regions, where not just developing countries but even the developed ones with some of the most advanced health systems are crippled and struggling to control the pandemic. Thus far, more than 500 million people have been infected, of which more than six (6) million have died (as of July 2022). Viral diversity, in particular amongst viruses of RNA genetic make-up, poses a significant challenge to the development of diagnostic, therapeutic and prophylactic interventions (Domingo and Holland 1997; Peck and Lauring 2018). Viral sequence variability within an infected individual can be an outcome of not just mutation, but also re-assortment and/or recombination, creating a diversity spectrum, termed as viral quasispecies (Domingo and Perales 2019), which can consist of one or more variants that are better fit for evolutionary selection. Sequence change, even that of a single amino acid substitution, can lead to immune escape and/or immunopathogenesis in some cases (Volkov et al. 2010).

The advances in genomics and proteomics approaches, coupled with the exponential reduction in the cost of sequencing, have allowed for a massive proliferation of sequence data. This provides unprecedented opportunities to study viral adaptation and evolution. Alignment-dependent approach is typically employed to study viral sequence divergence and conservation (Thompson et al. 2011). Conserved sequence regions can capture the genotypic diversity of majority or all historically reported variants of a virus, and likely that of future variants (Batzloff et al. 2004; Sitbon and Pietrokovski 2007; Haubold et al. 2011; Yang et al. 2015). Such regions are, however, limited in viral species that are highly variable, namely influenza A virus and human immunodeficiency virus 1 (HIV-1), amongst others, and even more so when applied to the search for universal vaccine targets that capture the diversity of multiple subtypes or subgroups of a highly diverse virus (Heiny et al. 2007; Hu et al. 2013). Naturally, aligning a large number of sequences of multiple viral species at the genus or family taxonomic lineage rank can become impractical, corresponding to a decline in the number and length of shared blocks of conserved regions that can anchor the alignment (Chong et al. 2021). Separately, multiple sequence alignment requires manual inspection to ascertain reliability, with correction of any misalignment. Further, aligning a large number of sequences can require a significant compute resource (Zielezinski et al. 2017; Ren et al. 2018). Therefore, there is a need for the development of alignment-free or -independent approaches to enable the study of viral sequence diversity at any rank of the taxonomic lineage.

The premise of an alignment-free approach is that it does not rely on the assignment of residue-residue correspondence to quantify sequence similarity. As such, the approach removes various restrictions, otherwise posed by an alignment-dependent approach for the study of sequence diversity (Ren et al. 2018). An alignment-free approach can involve performing an exhaustive search to determine the minimal set of sequences required to capture the diversity within a given dataset (Chong et al. 2021). The minimal set is herein defined as the smallest possible number of unique sequences required to represent the diversity inherent in the entire repertoire of overlapping *k*-mers encoded by all the unique sequences in a given dataset. The complete set of overlapping *k*-mers can be termed as part of the peptidome of the dataset, and thus the minimal set derived for a specific *k*-mer length is representative of the peptidome diversity relevant to the *k*-mer. Data compression of a given dataset to generate the minimal set is achieved at two levels. First, the redundant reduction (RR) of the dataset or the removal of duplicate sequences, which are common in public databases. Second, the non-redundant reduction (NRR) of the dataset, which is the removal of unique sequences whose entire repertoire of overlapping *k*-mer(s) can be represented by other unique sequences and thus, rendering them redundant to the collective pool of peptidome sequence diversity relevant to the *k*-mer. The compression can be significant and offers important insights into the effective sequence diversity and evolution of viruses when applied not just at the species level (such as analysis between specific proteins or proteome-wide), but at any rank of viral taxonomy lineage, such as genus, family or even at the highest, superkingdom level (all reported viruses). The study of minimal set has been previously reported for important viruses, such as dengue virus (Khan 2005; Khan et al. 2006) and influenza A virus (Tan 2009).

We have recently developed a novel algorithm for the search of a minimal set (Chong et al. 2021), which is improved and scalable for massive datasets, compared to the earlier iteration (Khan 2005; Khan et al. 2006). This has been implemented as a tool, UNIQmin to allow for a user-specific search for a minimal set of any *k*-mer at any rank of viral taxonomy lineage. The tool is publicly available via GitHub (https://github.com/ChongLC/MinimalSetofViralPeptidome-UNIQmin) and PyPI (https://pypi.org/project/uniqmin). The utility of the tool has been demonstrated for the sub-species *Severe acute respiratory syndrome coronavirus 2* (SARS-CoV-2), species *Dengue virus* and *Severe acute respiratory syndrome-related coronavirus*, genus *Flavivirus* and *Betacoronavirus*, family *Flaviviridae* and *Coronaviridae*, and even at the superkingdom rank, all reported *Viruses* (Chong et al. 2021; Chong and Khan 2022). Herein, we focus on describing a detailed three-part protocol (Figure 1) utilizing UNIQmin to address the issue of analysing viral diversity, alignment-free at any rank of the taxonomy lineage, which is much needed in this big data era for our future preparedness of a viral epidemic. The monkeypox virus (MPX) is used as a case study given the recent surge of human cases in countries where the disease is not typically reported. It is hoped that the detailed protocol will encourage broader application of the tool for alignment-free viral diversity studies.

**Figure 1.**
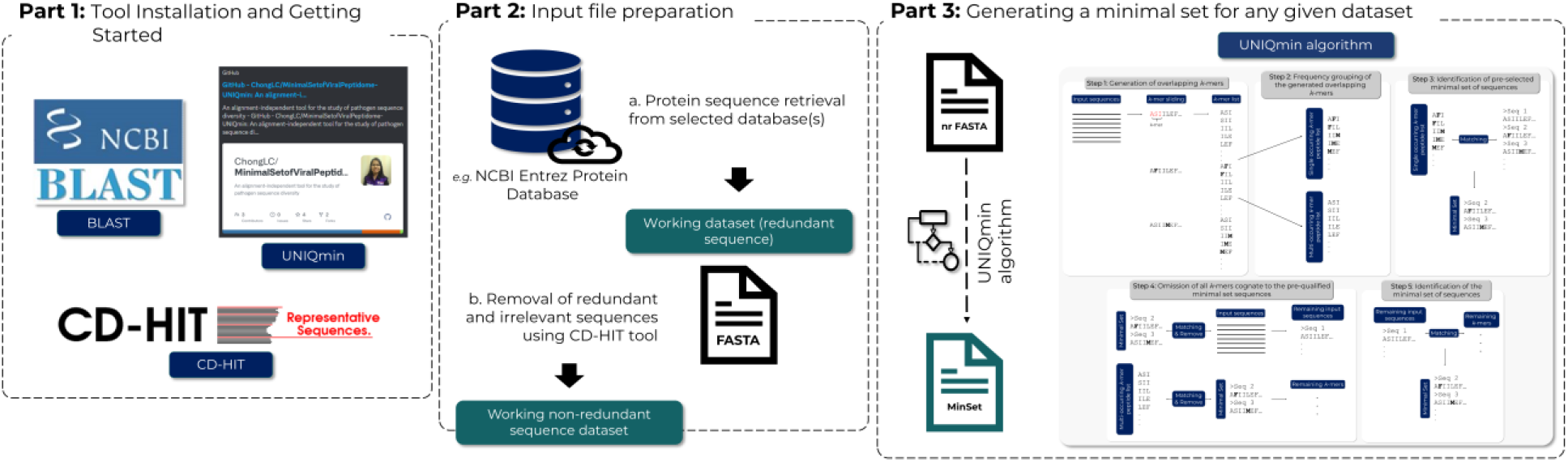
An overview of the protocol to study viral sequence diversity across taxonomic lineages by use of UNIQmin, an alignment-free tool. Protein sequences of interest are retrieved from a selected database as the working dataset. The dataset is removed of redundant and irrelevant sequences by use of CD-HIT, resulting in a non-redundant (nr) sequence dataset. UNIQmin is then applied to the nr dataset to generate a minimal set of sequences, which captures the peptidome diversity present within the nr dataset. Analysis of the minimal set can offer insights into the effective sequence diversity and evolution of viruses when applied across the taxonomy lineage.

## Protocol

### Part 1 — Tool Installation and Getting Started

The protocol presented here uses different software, all of which can be executed on a command-line interface (CLI) terminal. The software are listed below: -

i. **B**asic **L**ocal **A**lignment **S**earch **T**ool (BLAST)
ii. Clustering tool, **C**luster **D**atabase at **H**igh **I**dentity with **T**olerance (CD-HIT)
iii. UNIQmin

BLAST is available at https://ftp.ncbi.nlm.nih.gov/blast/executables/blast+/LATEST/ and a user can select according to the operating system. CD-HIT can be downloaded locally from http://weizhongli-lab.org/cd-hit/download.php. UNIQmin is publicly available at GitHub (https://github.com/ChongLC/MinimalSetofViralPeptidome-UNIQmin) and PyPI (https://pypi.org/project/uniqmin). UNIQmin is dependent on a few separate packages, namely pandas, biopython and pyahocorasick, and thus also requires them to be installed.

### Part 2 — Input File Preparation

Viral sequence data, a pre-requisite and key element for the study of diversity, is available in abundance in various public repositories, such as the **N**ational **C**enter for **B**iotechnology **I**nformation (NCBI; https://www.ncbi.nlm.nih.gov/) Entrez Protein (NR) database (Sayers et al. 2021). Derived or secondary sources, such as the NCBI Virus (https://www.ncbi.nlm.nih.gov/labs/virus/) (Hatcher et al. 2017) and **Vi**rus **P**athogen Database and Analysis **R**esource (ViPR; https://www.viprbrc.org/) (Pickett et al. 2012), among others, have also become commonly used resource for viral sequence data. They offer better data integration through other internal and extra sources, and provide internal curation, ease of data visualisation and download via various options for search result customisation. However, they can differ in their depth and breadth of curation, species coverage, and number of sequences. For example, ViPR covers 7,124 species/subgroups (as of January 2022), relative to NCBI Virus and NR, which in contrast cover approximately seven-fold more viral species/subgroups (47,823; as of January 2022); however, for a specific virus species that is covered by all the three databases, the NCBI Virus provides the best user-experience, in general.

Specialist databases are only available for a select few viruses, such as influenza A virus, human immunodeficiency virus (HIV-1) and the virus responsible for the current COVID-19 pandemic, SARS-CoV-2, among others, typically enriched with high curations (Stoesser et al. 2004; Khare et al. 2021). The **G**lobal **I**nitiative on **S**haring **A**ll **I**nfluenza **D**ata (GISAID; https://www.gisaid.org/), which started with EpiFlu™ database, later expanded to EpiCoV™ and EpiRSV™ and most recently added EpiPox™ boasts 1,651,325, 307,701,631, 24,420 and 482 sequence records (as of July 2022) for influenza viruses, SARS-CoV-2, respiratory syncytial virus (RSV) and monkeypox virus (MPX) respectively, with readily available metadata that facilitates “on-the-fly” analyses. These specialist databases are exemplary in their approach to biological data-warehousing (Schonbach et al. 2000; Neogi et al. 2013), such that they have become the preferred repository for community sequence submission, and in the case of GISAID, with numbers that far surpass the traditional primary repositories and even other specialist databases.

The wide availability, low-cost, and portability of next generation sequencing (NGS) offers unprecedented opportunity to the research community for sequencing of viruses of interest, including in real-time at the point/site of sample collection (nanopore technology). It is not uncommon for sequencing projects to output related or identical strains, which contribute to sampling bias (Koh et al. 2003). The removal of identical sequences may be necessary for a diversity study, to not only minimize the bias, but also reduce the demand on computational resources. In this protocol, we will showcase a standard workflow for data preparation.

#### 2.1. Retrieve viral protein sequences from publicly available database of choice

Sequence retrieval for a viral species of interest from the primary NCBI NR database is preferably done via the NCBI Taxonomy Browser, by searching for the species taxonomy identifier (txid) or the species name. It is important to ensure that the correct species has been determined from the search, as viruses can have similar names. For example, hepatitis A (HAV), hepatitis B (HBV), and hepatitis C (HCV) viruses, which appear related based on the common names have distinct lineages and thus, scientific names, such as species *Hepatovirus A* of the genus *Hepatovirus*, species *Hepatitis B* virus of the genus *Orthohepadnavirusvirus*, and species *Hepacivirus C* of the genus *Hepacivirus*, respectively. Thus, one should check the species lineage and cross-reference with the literature for the specific species of interest.

The species *Monkeypox virus* (MPX) is used as a case study to demonstrate data retrieval. Navigate the web client to the NCBI Taxonomy Browser (https://www.ncbi.nlm.nih.gov/Taxonomy/Browser/www.tax.cgi) (Figure 2) and search using the respective txid “10244”, followed by a download of the sequences in FASTA format.

**Figure 2.**
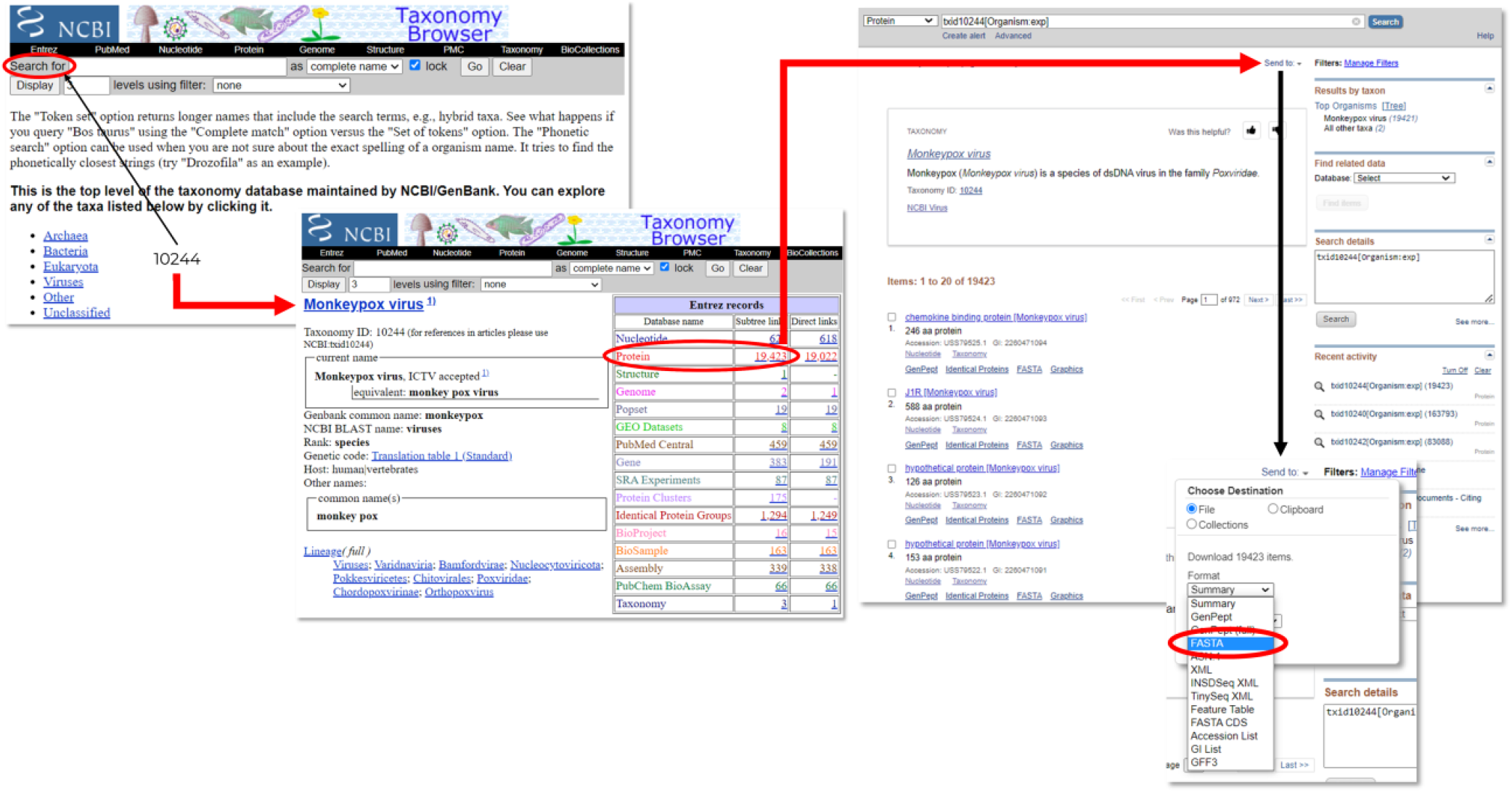
Data retrieval from the National Center for Biotechnology Information (NCBI) Entrez Protein (NR) Database for all reported *Monkeypox virus* protein sequences via the NCBI Taxonomy Browser, using the species taxonomy identifier (txid) “10244”.

Sequence data download should be accompanied with metadata download using the “GenPept (full)” format option. The metadata is useful for data processing, especially for filtering irrelevant sequences. The full records of nucleotide sequences should also be downloaded in ‘FASTA’ format and the metadata in “GenBank (full)” format. They can serve as a reference for comparative analysis.

It should be noted that since no alignment is to be done, both full-length and partial sequences can be included in the dataset for a comprehensive analysis of diversity. In an alignment-dependent study, partial sequences are a common source of spurious alignment, and hence it may be desired that they are filtered out from analyses, particularly when the number of such partial sequences may be prohibitively large, or the protein is of high diversity. Otherwise, in such cases, inspection and correction of misalignment can be a challenge and highly subjective. These concerns are not applicable for an alignment-free approach.

#### 2.2. [Optional] Extract selected viral protein(s) of interest from the retrieved data

A dataset retrieved from a primary or a secondary data resource(s) for a viral species of interest would comprise sequences of the proteins encoded by the viral genome. It may be in the interest of the user to analyse only one, a group, or all of the proteins encoded. Depending on the architecture of the virus genome, it is possible that a protein record would provide the sequence of only a protein encoded by the genome, as in the case of influenza virus (segmented genome) or provide the sequence of one, more than one, or all the proteins encoded by the genome, as in the case of dengue virus (polyprotein translation of the genome).

A BLAST (Altschul et al. 1990) search can be carried out to facilitate the extraction of the protein of interest. The retrieved dataset is used to construct a searchable database, whilst a reference sequence of the protein of interest can serve as the query for the BLASTp search. The reference sequence can be identified through a search in highly curated protein databases, such as UniProt (https://www.uniprot.org/) (Bateman et al. 2021) or RefSeq (https://www.ncbi.nlm.nih.gov/refseq/) (O’Leary et al. 2016).

#### 2.3. Remove redundant sequences from the dataset to be analysed

The tool, **C**luster **D**atabase at **H**igh **I**dentity with **T**olerance (CD-HIT; http://weizhong-lab.ucsd.edu/cd-hit/) is ideal for removal of redundant sequences at a given percentage similarity threshold of choice (Li and Godzik 2006). There are two ways to execute CD-HIT, either locally or using the webserver. One can load the sequence dataset to be analysed in FASTA format to the webserver (http://weizhong-lab.ucsd.edu/cdhit_suite/cgi-bin/index.cgi?cmd=cd-hit) and set the parameters. Removal of identical rather than similar sequences is preferred herein, and thus, the sequence identity cut-off should be set to 1, indicating 100% identity. Add desired email address for job checking and click on the ‘Submit’ button. Local client version of CD-HIT is preferred for large datasets. The clustering tool, **M**any-against-**M**any **seq**uence **s**earching (MMseqs2; https://github.com/soedinglab/MMseqs2) is an alternative to CD-HIT for big data, with an ultra-fast execution time (Steinegger and Söding 2018).

#### 2.4. Calculate the percentage of redundant reduction (RR) for the selected dataset analysed

Redundant reduction (RR) is defined as the percentage of identical sequences removed from the retrieved dataset and is calculated by use of the equation (Eq.) 1 below:

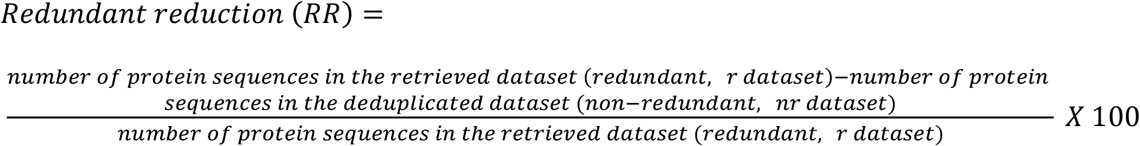

where r and nr datasets in Eq. 1 are referred to as the retrieved dataset containing redundant sequences and the dataset containing deduplicated sequences, respectively.

### Part 3 — Generation of minimal set for any given dataset using UNIQmin

This part demonstrates the generation of a minimal set for a dataset of interest. As per the definition of the minimal set, the *k*-mer size needs to be defined. Various *k*-mer sizes may be explored; see Chong *et al*. 2021 for the various considerations. Herein, as an example, we will utilise the *k*-mer size of nine (9-mer) for immunological applications, such as studying the viral diversity in the context of the cellular immune response (antigenic diversity). Peptides of different binding length can be recognised by the human leukocyte antigen (HLA) molecules. HLA-I molecules can bind peptides of length, ranging from eight (8) to 15 amino acids (aa), with nine (9) aa being the typical length; HLA class II molecules can bind longer peptides, up to 22 aa, with a binding core of nine (9) aa. The tool UNIQmin is used for the generation of the minimal set. The algorithm is detailed in Chong *et al*. 2021 and on the GitHub page (https://github.com/ChongLC/MinimalSetofViralPeptidome-UNIQmin), while the step-by-step procedure of the algorithm is provided in the README page of the ‘PythonScript’ folder (https://github.com/ChongLC/MinimalSetofViralPeptidome-UNIQmin/blob/master/PythonScript/README.md).

#### 3.1. Execute UNIQmin with the desired input file (such as “exampleinput.fas”)

The non-redundant (nr) file in FASTA format from Part 2 is used as the input. Execution of UNIQmin can be performed by use of the command below:

> *uniqmin -i exampleinput*.*fas -o example -k 9 -t 2*

Arguments:

*-i* *the path of the input file*
*-o* *the path of the output directory to be created*
*-k* *the length of the k-mers to be used*
*-t* *the number of threads to be used*

#### 3.2. Calculate the percentage of non-redundant reduction (NRR) for the generated minimal set

The generated minimal sequence set will be in the folder named as ‘minimal set’. NRR is defined as the percentage of non-redundant sequences removed from a deduplicated dataset and is calculated by the use of Eq. 2:

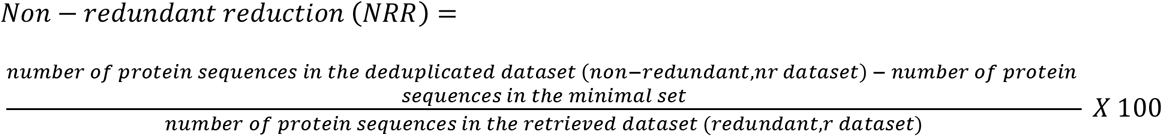

where r and nr datasets in Eq. 2 are referred to as the retrieved dataset containing redundant sequences and the dataset containing deduplicated sequences, respectively.

#### 3.3. Calculate the percentage of total reduction (TR) for the generated minimal set

TR is defined as the percentage of non-redundant sequences removed from a deduplicated dataset and is calculated using Eq. 3:

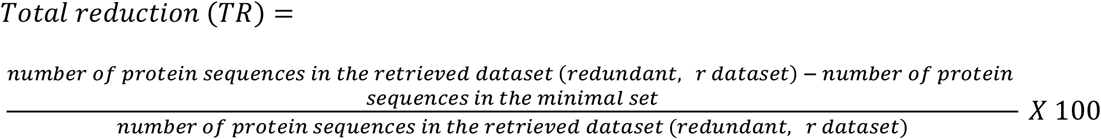

where r and nr datasets in Eq. 3 are referred to as the retrieved dataset containing redundant sequences and the dataset containing deduplicated sequences, respectively.

## Demo

### Comparative minimal set analysis across taxonomic lineage ranks: a case study of monkeypox virus

Viruses are classified based on similarity into groups, termed as taxa. Such classification enables a survey on the extent of virus genomic diversity, possibly leading to novel insights and future research on viral origin and evolutionary relationships (Simmonds et al. 2017). Starting from 2017, the hierarchy of virus taxonomy has been changed from a five-rank to 15-rank structure, including eight primary and seven derivative ranks. As of April 2022, the **I**nternational **C**ommittee on **T**axonomy of **V**iruses (ICTV) classified all known viruses into 10,434 species, 2,606 genera, 233 families, and 65 orders (https://talk.ictvonline.org/files/master-species-lists/).

Herein, we showcase the identification of minimal sets across higher taxonomic lineage ranks, by demonstrating for the species, genus and family levels of monkeypox virus (MPX). Such a comparative analysis, which would not possible via alignment-dependent approaches, provides for a broader understanding of minimal sets across the viral taxonomic lineage ranks, and in turn can offer a holistic understanding of sequence diversity across viral relations (from closer to distant).

1. Retrieve protein sequences of a selected virus from the publicly available database(s) of choice, across taxonomic lineage ranks (species, genus, and family ranks will be used as an example herein). Following the Part 2 of the protocol herein, all reported protein sequences of MPX across the higher lineage ranks (species, genus and family) were collected from the NCBI Entrez Protein (NR) Database, using the respective IDs (txid: 10244 for *Monkeypox virus* (species rank); 10242 for *Orthopoxvirus* (genus rank); and 10240 for *Poxviridae* (family rank)) via the NCBI Taxonomy Browser. The protein sequences in FASTA format, with the metadata in GenPept (full) format were downloaded for each rank. As of July 2022, a total of 19,423, 83,088, and 163,793 viral protein sequences of species *Monkeypox virus*, genus *Orthopoxvirus* and family *Poxviridae* were extracted, respectively.
2. Deduplicate the retrieved datasets. See Part 2.2 of the protocol. The retrieved dataset of 19,423 reported protein sequence of MPX was deduplicated by use of CD-HIT, resulting in a nr dataset of 1,245 sequences. Likewise, the deduplication of the retrieved dataset of the genus *Orthopoxvirus* (consisting of 83,088 protein sequences) resulted in an nr dataset of 15,523 sequences. Similarly, the retrieved sequences for the family *Poxviridae* were deduplicated, resulting in 34,782 nr sequences.
3. Implement UNIQmin to generate a minimal set of sequences for each dataset. See Part 3.1 of the protocol. The deduplicated dataset of MPX with 1,245 nr sequences was used as an input for UNIQmin, which generated a minimal set of 866 sequences. Similarly applying UNIQmin to the deduplicated datasets of the genus *Orthopoxvirus* and the family *Poxviridae* returned minimal sets of 9,146 and 25,618 sequences, respectively.
4. Calculate the percentage of RR, NRR, and TR for each of the minimal sets. See Parts 2.4, 3.2 and 3.3 of the protocol. Percentage of RR, NRR, and TR was calculated using Eqss. 1, 2, and 3, respectively. A summary of all the three reductions for MPX at the species, genus and family ranks is shown in Table 1 and Figure 3.
5. Compare and contrast the percentages of RR, NRR and TR for the selected virus across the taxonomic lineage ranks and analyse the relationship of all the reductions. A significant RR of ∼93.6% was observed at the species rank for MPX, while the NRR was only a mere ∼2.0%. Thus, less than 5% (866; TR of ∼95.5%) of the reported monkeypox protein sequences (19,423) are required to represent the inherent viral peptidome diversity at the species rank. The RR for MPX (species) was copious and is expected to vary from virus to virus. The NRR (∼2.0%) was relatively small and is expected to increase, transitioning from the species rank (the smallest dataset) to the family rank (the largest dataset) as the nr dataset increases to comprise more species from the different genera. Naturally, the TR is largely influenced by the RR. The RR at the genus rank was ∼81.3%, whereas the NRR was ∼7.7%, with ∼10.9% (9,146) of the reported genus sequences (83,088) required to represent the inherent viral peptidome diversity at the genus rank. The diversity was higher at the family rank: RR of ∼78.8% and NRR of ∼5.6%, with ∼15.6% (25,618) of the reported family sequences (163,793) required to represent the peptidome diversity at the rank. The peptidome diversity is limited at the species rank, which is favourable for the development of intervention strategies. A comparative analysis across different viruses at multiple lineage ranks can provide valuable insights in terms of trends of how the sequence increase at the redundant and non-redundant levels contribute to the effective diversity in the minimal set.

**Table 1.** A summary of all the three reductions (redundant (RR), non-redundant (NRR), and total (TR) reductions) across the taxonomic lineages ranks of monkeypox virus, namely species, genus, and family.

| Taxonomic lineage rank | Number of protein sequences<br>(r dataset \| nr dataset \| minimal<br>dataset) | Percentage of reduction<br>(%; 1 d.p.) |  |  |
| --- | --- | --- | --- | --- |
|  |  | RR | NRR | TR<sup>#</sup> |
| Species – *Monkeypox virus* * | 19,423 \| 1,245 \| 866 | 93.6 | 2.0 | 95.5 |
| Genus – *Orthopoxvirus* | 83,088 \| 15,523 \| 9,146 | 81.3 | 7.7 | 89.1 |
| Family – *Poxviridae* | 163,793 \| 34,782 \| 25,618 | 78.8 | 5.6 | 84.4 |

Abbreviation: \* MPX; <sup>#</sup> The addition of RR and NRR may not equal to TR due to rounding issue; r, redundant; nr, non-redundant; d.p., decimal place

**Figure 3.**
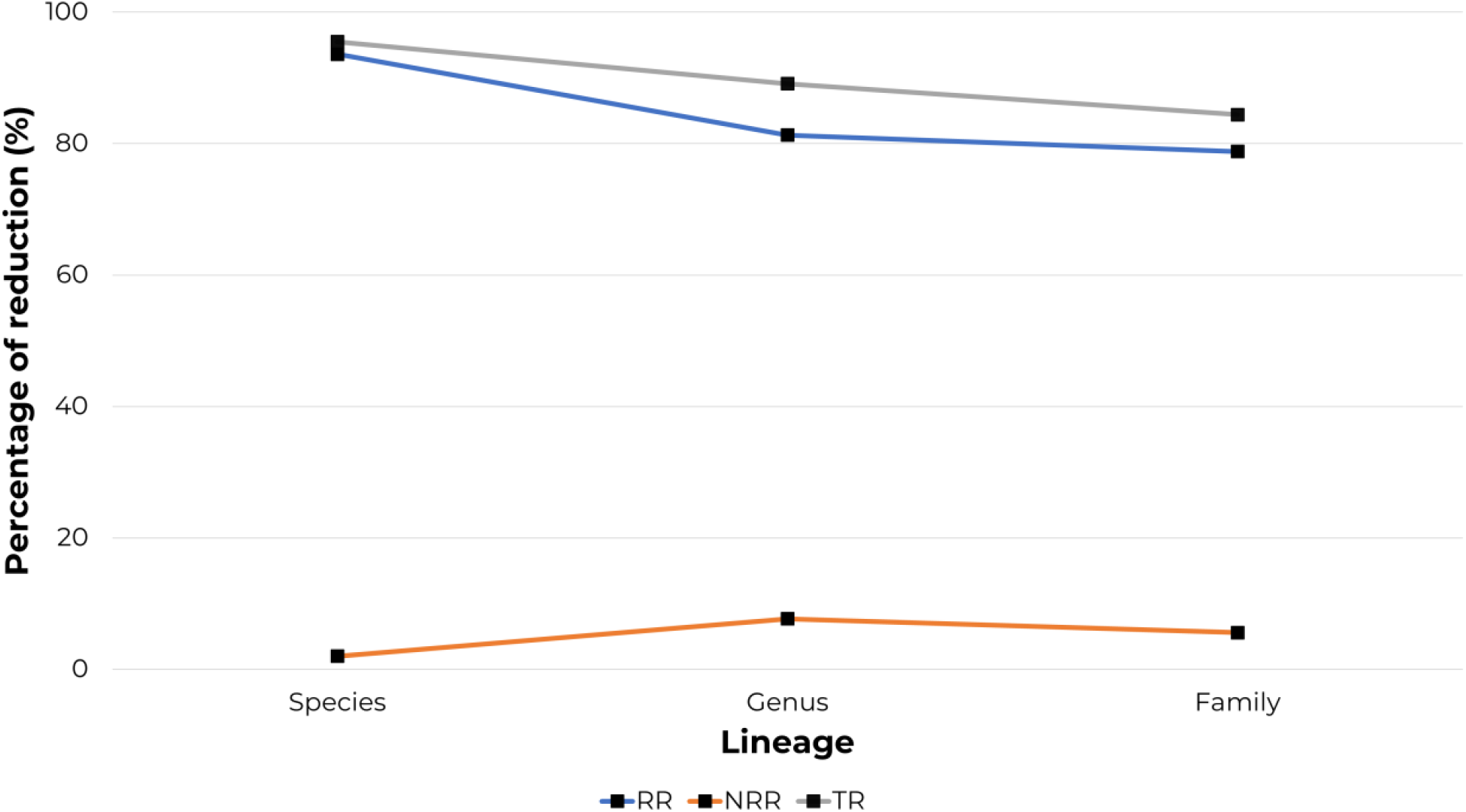
Percentage of reductions (%) for *Monkeypox virus* (MPX) across the viral taxonomic lineage ranks, namely species, genus, and family. Abbreviations: RR, redundant reduction, NRR, non-redundant reduction, TR, total reduction.

## Discussion

Viral diversity, an outcome of various evolutionary forces such as mutation, recombination, and re-assortment, plays a prominent role in disease emergence and control. UNIQmin enables study of viral diversity at the protein level in the context of peptidome degeneracies. The rapid surge in sequence data has necessitated alignment-free approaches to study diversity, in particular at higher taxonomic lineages. The concept of minimal set enables the study of sequence diversity without alignment, across taxonomic lineages. This is facilitated by the tool UNIQmin, which simply takes sequence data in FASTA format and outputs a compressed minimal set. Additionally, the compression offers a much-reduced data for downstream analyses without compromising on the peptidome diversity and is particularly beneficial in the case of big-data. Herein, we provided a detailed, systematic, step-by-step protocol on tool installation and getting started (Part 1), input file preparation (Part 2), and generation of minimal set for any given dataset using UNIQmin (Part 3). UNIQmin is applicable to viral and possibly other pathogenic microorganisms, with the possibility of dissecting the diversity spatio-temporally to allow for comparative analyses.

Herein, we also performed alignment-free study of monkeypox viral diversity (species level) and its higher taxonomic lineage ranks (genus and family) as a case study. Only less than 5% of the reported monkeypox protein sequences are required to represent the inherent viral peptidome diversity at the species rank, which increased to less than 16% at the family rank. The findings have important implications in the design of vaccines, drugs, and diagnostics, as only a small number of sequences are required for coverage of the MPX viral diversity.

## Data Availability

The tool and the data underlying this article are available in GitHub at https://github.com/ChongLC/MinimalSetofViralPeptidome-UNIQmin and https://github.com/ChongLC/UNIQmin_PublicationData, respectively. The datasets were derived from sources in the public domain, namely NCBI Protein database (https://www.ncbi.nlm.nih.gov/protein). The tool is also available at PyPI (https://pypi.org/project/uniqmin/).

## Key points

- UNIQmin offers an alignment-free approach for the study of viral sequence diversity at any given rank of taxonomy lineage and is big data ready
- The tool performs an exhaustive search to determine the minimal set of sequences required to capture the peptidome diversity within a given dataset.
- As an alignment-free approach removes various restrictions, otherwise posed by an alignment-dependent approach for the study of sequence diversity.
- The problem-solving protocol and case study described herein facilitate systematic diversity studies across viral taxonomic lineages.
- Alignment-free sequence diversity study of monkeypox virus across its taxonomy lineage (species, genus and family) revealed that only less than 5% of the reported monkeypox protein sequences are required to represent the inherent viral peptidome diversity at the species rank, which increased to ∼16% at the family rank. The low peptidome diversity, in particular at the species rank, is favourable for the development of intervention strategies.

## Acknowledgements

We gratefully acknowledge the support of Mr. Lim Wei Lun and Mr. Stephen Sugiharto in helping with the maintenance of UNIQmin.

